# Regular Architecture (RegArch): A standard expression language for describing protein architectures

**DOI:** 10.1101/679910

**Authors:** Davi R. Ortega, Grant J. Jensen

## Abstract

Domain architecture – the arrangement of features in a protein – exhibits syntactic patterns similar to the grammar of a language. This feature enables pattern mining for protein function prediction, comparative genomics, and studies of molecular evolution and complexity. To facilitate such work, here we propose Regular Architecture (RegArch), an expression language to describe syntactic patterns in protein architectures. Like the well-known Regular Expressions for text, RegArchs codify positional and non-positional patterns of elements into nested JSON objects. We describe the standard and provide a reference implementation in JavaScript to parse RegArchs and match annotated proteins.

## Introduction

Protein sequences can be compared to sentences in a natural language: each amino acid corresponding to a letter, and domains to words. Recently it was shown that, just as with language, evolutionary pressures on proteins’ biological roles give rise to a grammar structure in the protein universe (Yu *et al.*, 2019). In other words, only a subset of all possible arrangements of words (domains) form meaningful (useful) sentences (architectures). In the context of natural languages, grammar structures can be described by syntactic patterns (Chomsky, 1957), which can be used to mine large volumes of text for specific information. These syntactic patterns in regular languages are commonly codified as Regular Expressions (Kleene, 1956). We propose an analogous standard for describing syntactic patterns in proteins, enabling similar information mining. We call this standard Regular Architecture (RegArch). Here, we briefly describe the standard and give an example of its use.

## Concept

The goal of RegArch is to standardize flexible expressions that can be used to define, and therefore search, complex patterns of protein architecture. By architecture, we mean the arrangement of functional features/domains in a protein. Standard BLAST searches reveal all proteins that contain a particular domain, but RegArch allows one to search for proteins that contain multiple domains, in particular orders. RegArch does not identify domains in sequences, it just searches already-annotated sequences for specific domain patterns. Thus the protein sequences have to have already been annotated, for instance by bioinformatics resources such as PFAM (which identifies PFAM domains (Finn *et al.*, 2014)), DAS (which identifies transmembrane regions (Cserzo *et al.*, 2004)), SignalP (which identifies signal peptides (Armenteros *et al.*, 2019)), or some other method (Ulrich and Zhulin, 2014; UniProt Consortium, 2018).

Each RegArch expression is built of two types of patterns: positional and non-positional. Positional patterns match features that occur in a specific order. This order can occur anywhere in the sequence. As for Regular Expressions, RegArchs define the beginning and end of the protein sequence, so a user can also specify that the sequence must start and/or end with positional features. RegArch syntax contains a wild card, so a user can specify a pattern consisting of any combination of defined and undefined (i.e. any domain in the PFAM database) features. Finally, patterns can specify more than one requirement for a particular position, allowing a user, for instance, to exclude specific domains. Non-positional patterns do not require elements to appear in order, therefore allowing a user to search for a feature anywhere in the protein sequence, or to require that a feature be absent. Multiple positional and non-positional patterns can be combined in a single, intricate RegArch.

See the Supplementary Material for details of how to define, write and organize RegArch expressions. We have also designed a reference implementation in JavaScript to search for annotated proteins that match RegArchs. Details of this implementation can be found in the Supplementary Material.

## Usage example

Figure 1 illustrates an application of RegArchs to bacterial chemoreceptors, proteins that modulate signaling networks involved in the control of various cellular functions including motility (Porter *et al.*, 2011). In the PFAM database, bacterial chemoreceptors are characterized by an MCPsignal protein domain (Zhulin, 2001). Therefore, to find all chemoreceptors, we would simply define a RegArch consisting of a single non-positional pattern specifying the presence of one or more MCPsignal domains (Fig 1A). Of course this could also be accomplished with a simple BLAST search or PSI-BLAST. This may be perfectly fine for certain purposes, but because of gene duplication, domain swapping, and other evolutionary processes, this strategy is inefficient to find chemoreceptors with more specific domain architecture requirements. RegArch allows more detailed and specific patterns to be searched for as well. Chemoreceptors can be classified on the basis of their membrane topology (Zhulin, 2001), for instance. To select chemoreceptors of topological class I (crosses membrane twice with both termini in the cytoplasm and a periplasmic domain), we would define a RegArch with a positional pattern starting with a wild-card protein domain sandwiched by two transmembrane regions (TMs), followed anywhere in the protein by an MCPsignal PFAM domain (Fig 1B). Now imagine that we want to search for a very specific pattern: an initial TM, followed by any PFAM domain except Cache_1, followed by another TM, any number of PFAM domains, and ending with an MCPsignal PFAM domain. In addition, we want to exclude any protein with a PAS domain anywhere in the sequence. The resulting RegArch is shown in Fig 1C. The result of an example search for the RegArchs just described is shown in Fig 1D.

**Figure 1:**
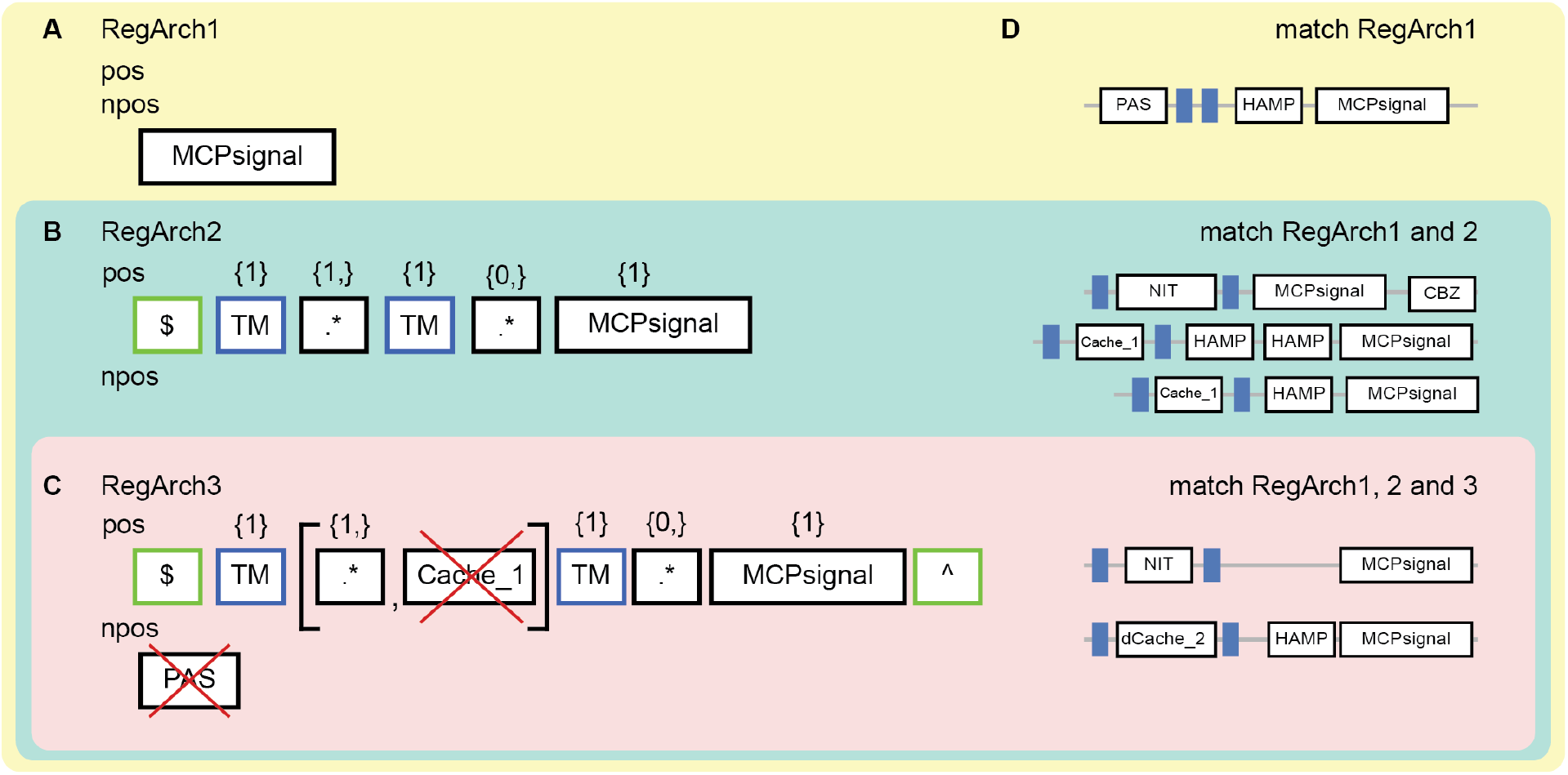
Example use of RegArchs to identify bacterial chemoreceptor proteins. (A-C) Three RegArchs of increasing stringency, described in the text, are shown. Note that RegArch1 contains only a non-positional pattern (npos), RegArch2 only a positional pattern (pos), and RegArch3 both. RegArch-specific features are denoted by a green border, PFAM features with black, and DAS features with blue. The symbols “$”, “^” and “.*” represent “start”, “end” and “any” respectively. D) RegArchs were used to search a set of annotated protein sequences with PFAM domains in white boxes with black borders and DAS predictions of TMs in blue. A sample of the protein feature architecture that matches the RegArchs is shown.

## Outlook

We expect RegArchs to improve automatic annotation of genomes, providing highly specific and efficient filters to identify multi-domain proteins with similar roles in biological pathways. RegArchs may also help reveal patterns in the grammar of protein architecture in different genomes, informing studies of molecular evolution. The quality of a dataset compiled by RegArchs strongly depends on the accuracy and completeness of underlying protein feature annotation. We therefore suggest tuning specificity, for instance by using highly specific patterns to build an initial high-confidence dataset that can later be used to search for related sequences. As the quality of computational predictions of protein features increases, so will the power of RegArch search algorithms.

## Supporting information

Supplementary Material: Manual

## Acknowledgments

The authors would like to thank the developers of MiST3: Luke Ulrich, Vadim Gumerov, and Ogun Adebali for helpful suggestions and comments on the manuscript. We also would like to thank Igor Zhulin for discussions on protein domain architectures that led to the idea of Regular Architecture. We also thank Dr. Catherine M. Oikonomou for helpful discussion and suggestions on the manuscript. This work was made possible through the support of the National Institutes of Health (grant R35 GM122588 to G.J.J.) and the John Templeton Foundation as part of the Boundaries of Life Initiative (grants 51250 & 60973 to G.J.J.).

