## Supplementary Material: Manual for "Regular Architecture (RegArch): A standard expression language for describing protein architectures"

### Regular Architecture

Regular Architecture is a sophisticated pattern matching tool to search for protein architectures. The name is partially borrowed from Regular Expressions.

This package evolved from [pfql] to a more straightforward API and a more robust code written in Typescript.

#### What RegArch is not

- RegArch is not an annotation tool.
- RegArch does not find patterns in protein sequences, it takes protein annotations as input.
- RegArch does not give partial matches

#### Specifications

- RegArch must be agnostic of annotation resources.
- RegArch must allow the use of annotations from multiple resources at the same time.
- RegArch must be able to deal with positional and non-positional pattern.

#### Building a pattern

`RegArch` uses JSON format to represent a pattern, in contrast with `RegExp` that uses strings.

`RegArch` patterns contain two sections: non-positional (`npos`) and positional (`pos`) and each of them contains an array of unit patterns.

##### A unit pattern

Because RegArch is agnostic of the resource used to annotate the protein sequence, we describe the name of the feature and the resource used. For example, to describe a Secretin domain from PFAM:

```
{
    name: "Secretin",
    resource: "pfam"
}
```

We can also include the number of times this domain should appear using `RegExp` notation with curly brackets `{}`. For example, to describe a pattern to match proteins with 2 or more `Secretin` domains:

```
{
    count: "{2,}",
    name: "Secretin",
```

```
    resource: "pfam"
}
```

#### Combining unit patterns

Unit patterns must be part of either positional or non-positional sections of the `RegArch` pattern. These sections are part of a JSON object with the key `patterns`. The following is a valid `RegArch` pattern:

```
{
    patterns: [
        {
            npos: [
                {
                    name: "CheW",
                    resource: "pfam"
                }
            ],
            pos: []
        }
    ]
}
```

This pattern matches all proteins that have 1 and only 1 CheW domain as defined in the PFAM domain database (resource), anywhere in the sequence. The `pos` keyword is mandatory even if no positional requirements are needed to describe this pattern.

Notice that the name of the feature and the resource are both conventions, but they must match how the proteins have been anottated (see `RegArch` annotation standards below).

It is also simple to write a positional pattern:

```
{
    patterns: [
        {
            npos: [],
            pos: [
                {
                    name: "CheW",
                    resource: "pfam"
                }
            ]
        }
    ]
}
```

This pattern matches the same proteins as the previous pattern: 1 and only 1 CheW anywhere in the sequence. That is because RegArch starts to require positional matches from the position it finds the first match.

To match proteins that start with a particular feature, we can rely on `RegArch` special features. The symbols `^` and `$` are both RegArch features meaning *start* of the sequence and *end* of the sequence, respectively. To only select proteins with 1 and only 1 CheW and no other feature what so ever, we must make a positional pattern:

```
{
    patterns: [
        {
            npos: [],
            pos: [
                {
                    name: "^",
                    resource: "regarch"
                },
                {
                    name: "CheW",
                    resource: "pfam"
                },
                {
                    name: "$",
                    resource: "regarch"
                },
            ]
        }
    ]
}
```

Notice that the pattern starts with the `^` from the resource `regarch` and ends with `$` from the resource `regarch`.

#### Complex combinations of patterns

We also can combine positional and non-positional rules in the same pattern:

```
{
    patterns: [
        {
            npos: [
                {
                    name: "HATPase_c",
                    resource: "pfam"
                },
            ],
            pos: [
                {
                    name: "CheW",
                    resource: "pfam"
                },
                {
                    name: "$",
                    resource: "regarch"
                },
            ]
```

```
        }
    ]
}
```

This pattern matches proteins that end with CheW domain and an HATPase_c anywhere in the sequence. Those are most likely histidine kinases from the chemotaxis pathway.

We can easily combine features from multiple resources. For example to match proteins that start with a transmembrane region predicted by DAS and ends with the MCPsignal domain from PFAM database:

```
{
    patterns: [
        {
            npos: [],
            pos: [
                {
                    name: "^",
                    resource: "regarch"
                },
                {
                    name: "TM",
                    resource: "das"
                },
                {
                    name: "MCPsignal",
                    resource: "pfam"
                },
                {
                    name: "$",
                    resource: "regarch"
                },
            ]
        }
    ]
}
```

Notice that `TM` was just a convention we made. DAS only annotate transmembrane regions and does not have a feature name for it. We called it as `TM` when parsing the DAS results and formatted the protein annotation as required by `RegArch`.

#### Logical AND and ORs

RegArch allows combining patterns in AND and OR logical operations. All unit patterns in non-positional arrays are interpreted as AND, meaning they all must match for the protein to be a match to the pattern. For example, a pattern to match every protein that has a CheW domain **AND** a Response_reg domain.

```
{
    patterns: [
        {
            npos: [
```

```
                {
                    name: "CheW",
                    resource: "pfam"
                },
                {
                    name: "Response_reg",
                    resource: "pfam"
                },
            ],
            pos: []
        }
    ]
}
```

To combine non-positional patterns with OR logical operator, we need to add a pattern object in the array. For example, to match proteins with CheW, Response_reg, **OR** both:

```
{
    patterns: [
        {
            npos: [
                {
                    name: "CheW",
                    resource: "pfam"
                }
            ],
            pos: []
        },
        {
            npos: [
                {
                    name: "Response_reg",
                    resource: "pfam"
                },
            ],
            pos: []
        }
    ]
}
```

**OR** operations are similar for positional patterns; we must repeat the pattern as a new element for the array `patterns` and change only the part we want.

We can also build positional patterns that require that multiple unit patterns match the same position. For that, `RegArch` positional patterns can be nested.

For example, to match proteins that have 1 occurrence of any PFAM domains except Cache_1 between two transmembranes. Thus, we must request that at the position between the two transmembrane regions two unit patterns must match at the same time: any pfam domain AND a {0} count for Cache_1 domain:

```
{
    patterns: [
```

```
        {
            npos: [],
            pos: [
                {
                    name: 'TM',
                    resource: 'das',
                },
                [
                    {
                        name: '.*',
                        resource: '.*',
                    },
                    {
                        count: '{0}',
                        name: 'Cache_1',
                        resource: 'pfam',
                    },
                ],
                {
                    name: 'TM',
                    resource: 'das',
                },
            ]
        }
    ]
}
```

This flexible standard allows for simple and complex patterns to be built and can help to compose protein sequence datasets efficiently.

### Formatting protein annotations

As with the patterns, `RegArch` also uses JSON formatted objects to represent protein annotations. The idea behind is that most bioinformatics study dealing with different types of information work with nested JSON objects where different keys representing different information for a single protein sequence. For this reason, `RegArch` considers the protein annotation as the value of the object with key `pa`.

The following is a valid `RegArch` protein annotation:

```
{
    "pa": {
        "pfam":[
            ["CheW",171,314,"..",0.1,2,137,"..",170,315,"..",113.1,3e-
36,4.3e-33,0.94],
            ["CheW",2,139,"..",0.1,4,136,"..",1,141,"[.",83.2,4.9e-27,7e-
24,0.93]
        ],
        "smart":[
            ["SM00260",2,137,4.9e-17],
            ["SM00260",166,311,1.5e-31]
        ]
    }
```

```
}
```

The information from each resource must appear in an array under the key with the resource's name. Each feature must appear as an array with the following format `["name_of_the_feature", start, stop]`. `RegArch` ignores other information in the rest of the array.

**These names (feature and resources) are the names that must match with the ones used in the patterns.**

### Scope of resources

Positional patterns must make sure that the next unit pattern matches the next protein annotation. However, because RegArch accepts annotation from multiple resources, it might be the case that some of the annotations are irrelevant for this particular pattern. For example, if we annotate a CheW protein with PFAM and SMART domains, we get something that looks like this:

```
{
    "pa": {
        "pfam":[["CheW",3,136,"..",4,2,135,"..",2,139,"..",115.7,1.3e-
37,6.6e-34,0.98]],
        "smart":[["SM00260",1,135,2.9e-32]]
        }
    },
}
```

If `RegArch` ignored the scope of resources, the following pattern would not match the above annotation:

```
{
    patterns: [
        {
            npos: [],
            pos: [
                {
                    name: "^",
                    resource: "regarch"
                },
                {
                    name: "CheW",
                    resource: "pfam"
                },
                {
                    name: "$",
                    resource: "regarch"
                },
            ]
        }
    ]
}
```

However, since there is only mention of the `pfam` resource, `RegArch` ignores the annotations of all other resources.

We can make a more stringent search for CheW proteins from both resources by running two `RegArch`, one for each resource, and requesting that both must be a match.

```
import { RegArch } from 'regarch'

const raPfam = new RegArch(patternUsingPfam)
const raSmart = new RegArch(patternUsingSmart)

const matchPfam = raPfam.exec(listOfAnnotatedProteins)
const matchSmart = raSmart.exec(listOfAnnotatedProteins)

const matchBoth = listOfAnnotatedProteins.filter((pa, i) => {
    return matchPfam[i] && matchSmart[i]
})
```

### The API

Much like `RegExp`, we instantiate a new instance of `RegArch` with a pattern. The only exposed method in a `RegArch` instance is `exec()`, which takes an array with protein annotations as input.

Pretty much like this:

```
import { RegArch } from 'regarch'

const ra = new RegArch(pattern)
const results = ra.exec(listOfAnnotatedProteins)
```

The `results` is a boolean array with true if the protein matches the pattern and false if it does not.

---

Now we have everything we need to use RegArch.

Happy matching!
